# TLR4 deficiency does not alter glaucomatous progression in a mouse model of chronic glaucoma

**DOI:** 10.1101/2024.06.07.597951

**Authors:** Chi Zhang, Marina Simón, Jeffrey M. Harder, Haeyn Lim, Christa Montgomery, Qing Wang, Simon W.M. John

**Affiliations:** Department of Ophthalmology, Columbia University Irving Medical Center, New York, NY; Mortimer B. Zuckerman Mind Brain Behavior Institute, Columbia University, New York, NY; The Jackson Laboratory, Bar Harbor, ME

## Abstract

Glaucoma is a leading cause of irreversible blindness worldwide. Toll-like receptor 4 (TLR4) is a pattern-recognition transmembrane receptor that induces neuroinflammatory processes in response to injury. *Tlr4* is highly expressed in ocular tissues and is known to modulate inflammatory processes in both anterior and posterior segment tissues. TLR4 activation can lead to mitochondrial dysfunction and metabolic deficits in inflammatory disorders. Due to its effects on inflammation and metabolism, TLR4 is a candidate to participate in glaucoma pathogenesis. It has been suggested as a therapeutic target based on studies using acute models, such as experimentally raising IOP to ischemia-inducing levels. Nevertheless, its role in chronic glaucoma needs further evaluation. In the current study, we investigated the role of TLR4 in an inherited mouse model of chronic glaucoma, DBA/2J. To do this, we analyzed the effect of *Tlr4* knockout (*Tlr4*^−/−^) on glaucoma-associated phenotypes in DBA/2J mice. Our studies found no significant differences in intraocular pressure, iris disease, or glaucomatous progression in *Tlr4*^−/−^ compared to *Tlr4*^+/+^ DBA/2J mice. These data do not identify a role for TLR4 in this chronic glaucoma, but further research is warranted to understand its role in other glaucoma models and different genetic contexts.

## Introduction

Glaucoma refers to a complex group of multifactorial diseases characterized by the progressive degeneration of retinal ganglion cells (RGCs) and visual field deficits^1,2^. Elevated intraocular pressure (IOP), genetics, and advanced age are considered the most important risk factors for glaucoma^2,3^. Both neuroinflammation and perturbed metabolism are implicated in the disease. Nevertheless, the mechanisms that lead to neural demise and degeneration in glaucoma are yet to be fully elucidated.

Neuroinflammation is involved in glaucomatous progression^1,2,4–8^. We have previously investigated the processes implicated in the initiation of glaucomatous damage and identified monocyte cell entry into the optic nerve head as an important pathogenic event in DBA/2J glaucoma^4,6,9,10^. Subsequently, macrophages were found to infiltrate axon bundles in the optic nerves of glaucoma patients^11^. Additionally, we and others have shown that mitochondrial dysfunction and impaired metabolism contribute to the development and progression of glaucoma^8,12–23^. In a vicious cycle, pro-inflammatory mediators (released by e.g., infiltrating macrophages) can trigger mitochondrial damage and amplify the neurodegeneration driven by mitochondrial dysfunction^17,24^.

Toll-like receptors (TLR) are a family of pattern-recognition transmembrane receptors and a centerpiece of the innate immune response. They can recognize external pathogen-associated molecular patterns (PAMPs) like bacterial lipopolysaccharide (LPS), as well as endogenous damage-associated molecular patterns (DAMPs). They play a significant role in inflammatory conditions and neurodegenerative diseases^25^. TLR4, in particular, is highly expressed in cells of the central nervous system and the retina, and its activation triggers multiple signaling cascades that lead to the release of inflammatory cytokines and immune modulators^26,27^. TLR4 has been implicated in the regulation of glucolipid metabolism^28^ and in mitochondrial dysfunction after LPS-induced TLR4 activation^29^. Additionally, TLR4 activation can induce metabolic changes in macrophages via mitochondrial reprogramming^30^.

TLR4 has been studied as a therapeutic target in preclinical models of eye disorders. Its activation has been linked to ocular inflammation, retinal damage following acute experimental elevation of IOP to an ischemic level, and retinal/ischemia-reperfusion injury^31–33^. In murine optic nerve crush, an acute model with relevance to glaucoma due to direct axon injury, blocking TLR4 suppressed inflammatory reactions and RGC loss^34,35^. In human tissues, increased expression of TLR4 was reported in glaucomatous human trabecular meshwork (TM), a tissue relevant to IOP elevation, and in glaucomatous human optic nerve head^36–38^. Finally, polymorphisms in *TLR4* are suggested to be involved in normal tension glaucoma and primary open-angle glaucoma (POAG), but further data are needed^39–42^.

Given the important effects of TLR4 on both metabolism and neuroinflammation, we functionally tested the role of TLR4 in a chronic glaucoma by comparing *Tlr4*-deficient (*Tlr4*^−/−^) and wild-type (*Tlr4*^*+/+*^) littermate mice on a DBA/2J background. DBA/2J mice are a model of hereditary glaucoma and are widely used for glaucoma research. Mutations in two genes impacting melanosomes cause pigment-dispersing iris disease, IOP elevation, and glaucoma in DBA/2J mice^43^. Variants in *PMEL*, a gene with a role in melanosome biology, cause pigmentary glaucoma in humans^44^. Thus, the initiating melanosomal etiology of DBA/2J glaucoma is similar to at least a subset of human pigmentary glaucoma. Importantly, findings in DBA/2J have been extended to POAG, the most common form of glaucoma, with initial promising outcomes in clinical trials^12,15,45,46^. In our experiments, we studied whether TLR4 plays a role in the development of DBA/2J glaucoma. We observed no significant differences in IOP, nerve damage, and other glaucoma-associated phenotypes between *Tlr4*^*+/+*^ and *Tlr4*^−/−^ DBA/2J mice. Our data indicate that TLR4 is not necessary for disease progression in this chronic glaucoma model.

## Materials and methods

### Mice

B6.B10ScN-*Tlr4*^*lps-del*^/JthJ mice (strain #007227) were obtained from The Jackson Laboratory^47,48^. These mice are *Tlr4*-deficient due to lack of exon 3 in the *Tlr4* gene. The *Tlr4* mutation was backcrossed to strain DBA/2J (strain #000671) for >10 generations to produce congenic mice with the *Tlr4*-deficient allele on the DBA/2J background (all experimental mice were ≥N10). For this study, congenic DBA/2J *Tlr4* heterozygote mice were intercrossed to produce *Tlr4* wild-type and homozygous mutant littermates. Both genotypes were housed together and analyzed simultaneously. All mice were housed in a 21°C environment with a 14-h light and 10-h dark cycle, fed with a 6% fat diet, and acidified (pH 2.8–3.2) drinking water^49^. Both female and male mice were used for analysis.

### Clinical slit-lamp examination

Mice underwent regular examinations using a slit lamp bio-microscope at 40X magnification throughout their lifespan, as described in previous studies^43,50,51^. The analysis started at 4 months of age, with subsequent examinations at 6, 8, 10, and 12 months. Phenotypic assessment of iris disease included the evaluation of iris atrophy, pigment dispersion, and transillumination. Mice were examined without anesthesia using standard handling, and all photographs were captured with identical camera settings.

### IOP measurement

IOP was assessed using the microneedle method as previously outlined in detail^52,53^. In brief, mice were anesthetized with an intraperitoneal injection of a combination of ketamine (99 mg/kg; Ketlar, Parke-Davis, Paramus, NJ, USA) and xylazine (9 mg/kg; Rompun, Phoenix Pharmaceutical, St Joseph, MO, USA) immediately before the IOP measurement. All IOP measurements for both genotypes were taken at the same time of the day.

### Nerve staining and evaluation of damage

The intracranial segments of the optic nerves were fixed in 0.8% paraformaldehyde, 1.22% glutaraldehyde, and 0.08 M Phosphate Buffer pH 7.4 at 4°C for 12 hours, followed by processing and embedding in plastic. The retro-orbital ends of the nerves were cut into 1 μm thick sections and stained with paraphenylenediamine (PPD) as previously described^54–56^. PPD stains the myelin sheath of all axons, while specifically darkly staining the axoplasm of damaged axons^56^. At least 3 sections were examined to determine the level of damage for each nerve. The optic nerves were determined to have 3 levels of damage as previously reported and validated against axon counting^9,12^: 1. No or early damage (NOE) - less than 5% of axons were damaged, no gliosis, indistinguishable from no glaucoma controls. These nerves have no damage by conventional criteria but are called no or early as some of them have early molecular changes that precede neurodegeneration. 2. Moderate damage (MOD) - had an average of 30% axonal loss and early gliosis. 3. Severe damage (SEV) - had more than 50% axonal loss and damage with prominent gliosis^4,10,23,54^. All nerves were evaluated by at least two masked investigators. In cases where the two investigators did not agree on the damage level a third investigator (also masked) analyzed the nerve and the most assigned damage level was used^54^.

### Retinal wholemounts Nissl staining and cell counting

Retinas from 12-month-old DBA/2J mice were Nissl-stained with cresyl violet as previously reported^57^. In short, eyes were fixed in 4% paraformaldehyde in 0.1M Phosphate Buffer pH 7.4 overnight at 4°C and then transferred into 0.4% paraformaldehyde in 0.1M Phosphate Buffer pH 7.4. Whole retinas were dissected from the eye, processed with 0.3% Triton X-100 and 3% hydrogen peroxide, flat-mounted onto glass slides, and stained for 1h in 1% cresyl violet in distilled water before being differentiated in 95% alcohol, 100% alcohol, and xylene. Eight 40x brightfield images were obtained (two per quadrant) of peripheral retina, equidistant from the peripheral edge. RGC layer cells were manually counted (endothelial cells excluded) and averaged across all eight images per retina.

### Statistical analysis

IOP measurements were compared between *Tlr4*^*+/+*^ and *Tlr4*^−/−^ mice at all ages with a two-way ANOVA. The level of nerve damage between *Tlr4*^*+/+*^ and *Tlr4*^−/−^ mice at each age was compared using Fisher’s exact test. RGC numbers were compared with a one-way ANOVA with post hoc Tukey’s HSD test.

## Results

### *Tlr4* mutation has no effect on pigment-dispersing iris disease or IOP

DBA/2J mice develop a pigment-dispersing iris disease that results in pigment accumulation in the ocular drainage tissues and subsequent IOP elevation. There were no differences in the onset or progression of the iris disease between mice of each *Tlr4* genotype (**Figure 1A**). At all ages (analyzed between 4 and 12 months old), both groups followed the usual anterior segment disease progression that we and others have previously characterized for DBA/2J mice^51,58–61^. DBA/2J mice typically have clear disease by 5 months of age, consisting of transillumination defects that involve progressive depigmentation and iris atrophy^51,58,59^. At 8 months of age, both *Tlr4*^*+/+*^ and *Tlr4*^−/−^ mice presented peripupillary accumulation of pigment, dispersed pigment, and transillumination defects. The depigmenting iris disease worsened by 12 months of age, with severe transillumination defects and iris atrophy in both groups. In the DBA/2J model, the IOP is increased in many mice by 8-9 months of age and starts declining after 12-13 months due to atrophy of the ciliary body^51,59^. We did not detect any differences in IOP between *Tlr4*^*+/+*^ and *Tlr4*^−/−^ mice (i.e., no effect of the genotype, p-value = 0.702, two-way ANOVA. **Figure 1B**).

**Figure 1.**
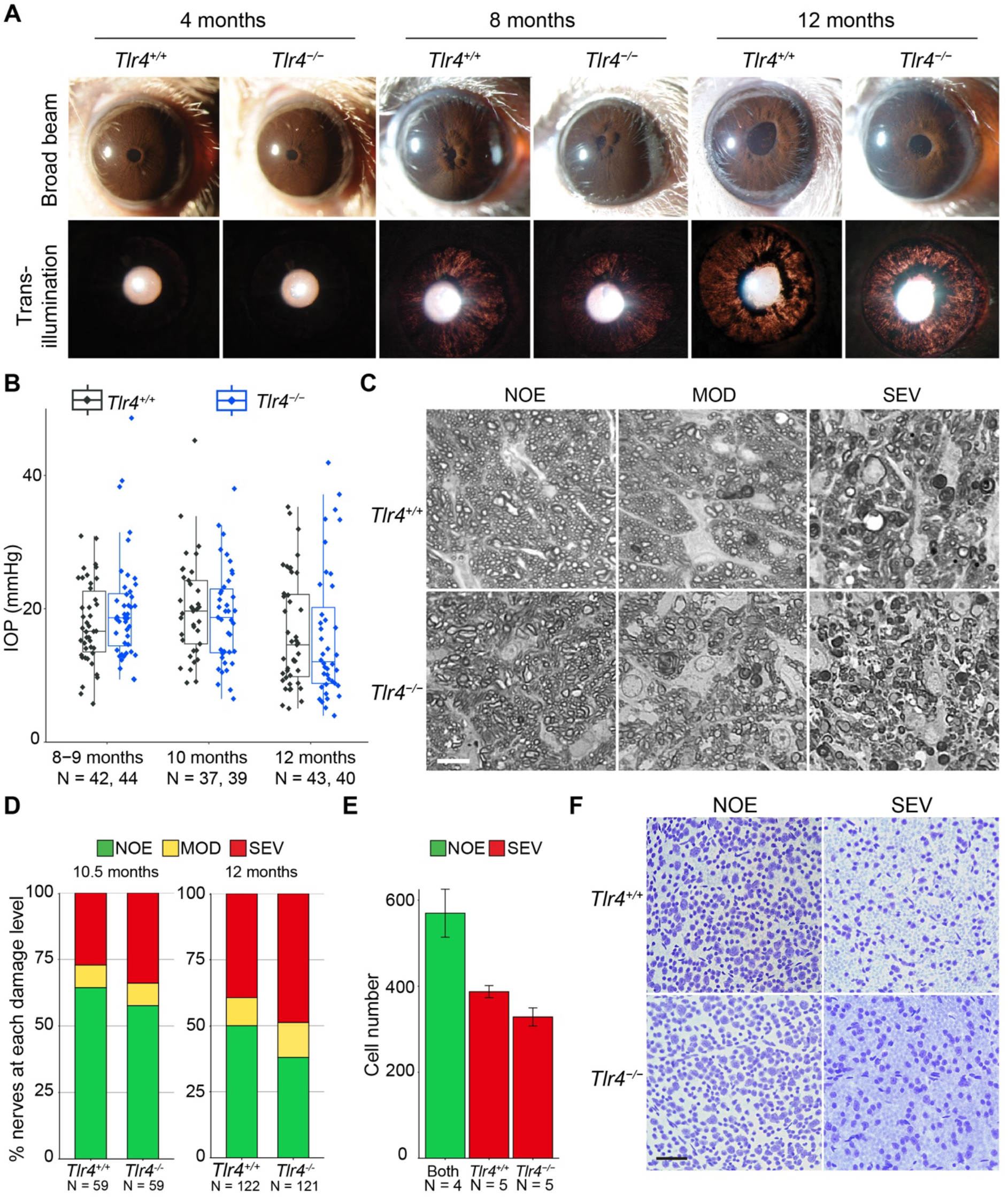
*Tlr4* deficiency does not influence the glaucoma phenotype of DBA/2J mice. **A**. Representative slit-lamp images of *Tlr4*^+/+^ (wild-type DBA/2J) and *Tlr4*^−/−^ eyes at 4, 8, and 12 months. The top row shows broad beam illumination, and the bottom row shows transillumination. The glaucoma-related changes in *Tlr4*^−/−^ mice developed at the same time as in *Tlr4*^*+/+*^ mice. At 4 months of age, iris disease is not yet evident in either of the groups. Eight-month-old *Tlr4*^+/+^ and Tlr4^−/−^ eyes present dispersed pigment and transillumination defects, which become more severe at 12 months of age. N > 20 mice per genotype at each age. **B**. IOP distributions at key ages. There was no significant effect of the genotype on IOP levels at any age (two-way ANOVA; *P* = 0.702). **C**. Representative images of nerves showing no/early (NOE), moderate (MOD), and severe (SEV) damage. Scale bar = 10 μm. **D**.Frequency distribution of optic nerve damage at 10.5 and 12 months of age. No significant differences were observed between the two genotypes of mice at any age (Fisher’s test at 10.5 months, *P* = 0.749; *P* = 0.169 at 12 months of age). **E**. RGC layer cell counts for eyes with no glaucoma (NOE, methods) or severe (SEV) glaucoma based on optic nerve damage. No significant differences in RGC layer cell numbers were observed between eyes of either genotype with severe glaucoma indicating that somal and axonal damage are not uncoupled by the *Tlr4* mutation (one-way ANOVA and post hoc Tukey’s HSD test, *Tlr4*^*+/+*^ SEV vs. *Tlr4*^−/−^ SEV; *P* = 0.384). **F**. Representative images of the retinas of *Tlr4*^*+/+*^ and *Tlr4*^−/−^ mice with no or severe nerve damage at 40X. Scale bar = 50 μm.

### *Tlr4* knockout has no effect on RGC numbers or axonal degeneration

To study the role of TLR4 in glaucomatous degeneration, we assessed the effects of *Tlr4* deficiency on optic nerve damage using PPD-stained nerve cross sections. We evaluated nerves at 10.5 and 12 months of age, two key glaucoma points frequently used for assessing glaucomatous neurodegeneration in this model^4,62^ (**Figure 1C-D**). There were no significant differences in nerve damage between genotypes at the two ages analyzed. Because the *Tlr4* mutant mice initially had a slightly higher incidence of severe damage at each age, we analyzed an even larger number of 12-month-old eyes. Despite the larger number of nerves, the difference remained statistically insignificant (Fisher’s test comparing *Tlr4*^*+/+*^ and *Tlr4*^−/−^ mice at 10.5 months of age, *P* = 0.749; *P* = 0.169 at 12 months of age). At 10.5 months of age, 27% of the *Tlr4*^*+/+*^ nerves had severe nerve damage (> 50% axonal loss), and this proportion increased to 39% at 12 months of age. For the *Tlr4*^−/−^ mice, 33.9% of the nerves had severe damage at 10.5 months of age, which increased to 48.8% by 12 months.

To evaluate potential axonal-somal uncoupling, where RGC somal survival is observed despite axon loss^55,57,63^, we analyzed RGC layer cell numbers in Nissl-stained flat-mounted retinas. To assess uncoupling, we counted and compared retinas of *Tlr4*^*+/+*^ and *Tlr4*^−/−^ eyes that had severe or no glaucomatous damage (**Figure 1E-F**). As expected, retinal cell counts in *Tlr4*^*+/+*^ and *Tlr4*^−/−^ mice with severe nerve damage were considerably reduced, but there was no significant effect of the genotype on this loss (*P* = 0.384 when comparing *Tlr4*^*+/+*^ and *Tlr4*^−/−^ retinas with severe damage, one-way ANOVA and post hoc Tukey’s HSD test). Thus, knocking out *Tlr4* does not prevent RGC death or glaucomatous nerve damage.

## Discussion

TLR4 targeting strategies have been studied in multiple models of diseases characterized by an inflammatory microenvironment^27,64^. TLR4 is expressed by a variety of ocular tissues and plays an important role in inflammatory eye diseases^65–67^. A study in mice showed that TLR4 contributes to retinal ischemia/reperfusion injury via NF-kB signaling, increasing the expression of proinflammatory genes^32^. Knockout of *Tlr4* led to a suppressed inflammatory response in the retina. Another study analyzed TLR4 downstream mechanisms and showed that TLR4 signaling mediated by caspase-8 controlled the production of the inflammatory cytokine interleukin 1β (IL-1β)^31^. Inhibition of either TLR4 or caspase-8 blocked the production of IL-1β and attenuated retinal ischemic damage after acute experimental IOP elevation. *Tlr4*^−/−^ mice have also been shown to be more resistant to optic nerve crush injury than wild-type mice^34^.

Growing evidence suggests that mitochondrial and metabolic disturbances with bioenergetic insufficiency contribute to the demise of RGCs in glaucoma^15,68–70^, while an extensive literature implicates neuroinflammatory processes^1,2,4,5^. TLR4 plays a role in multiple metabolic processes, affects mitochondrial function in inflammatory diseases, and has been proposed to have a role in acute glaucoma based on an induced, acute model with very high IOP elevation (ischemia-inducing)^31^, but a role in chronic glaucoma is not assessed. Thus, we determined its role in the chronic DBA/2J glaucoma model. Our experiments did not detect any differences in the glaucoma-related phenotype between *Tlr4*^−/−^ and *Tlr4*^+/+^ DBA/2J mice.

Both genotypes developed comparable levels of IOP elevation, RGC loss, and optic nerve damage. Our data do not provide evidence for a role of TLR4 in this chronic glaucoma.

The difference between our results and those in more acute models is not surprising as different types and degrees of insult can induce differing damaging mechanisms. The previous study in the acute model used an IOP of 110 mmHg^31^, and such a degree of IOP elevation does not occur in chronic glaucoma or even in most human patients with acute glaucoma. Additionally, optic nerve crush is meant to model direct axon injury in glaucoma, but there are differences to IOP-induced glaucoma including transcriptomic differences^71,72^.

Our studies indicate that the lack of *Tlr4* does not influence glaucoma progression in DBA/2J glaucoma under our standard housing conditions. It is well known that multiple factors influence disease progression, such as genetic background and environment^73–75^. Various factors may influence the effects of TLR4 on glaucoma. TLR4 mediates inflammation in response to infection and endogenous molecules such as saturated fatty acids^76,77^. Thus, the presence of specific infectious or commensal microbes or the nature of diet, including saturated fat content, may impact its role in glaucoma. The specific mutation is also expected to play a role^75^. Further experiments are needed to understand the role of TLR4 in other chronic glaucoma models and in different microbial and dietary environments.

## Declarations

### Ethics approval

All mice were treated in accordance with the Association for Research in Vision and Ophthalmology’s statement on the use of animals in ophthalmic research. All animal procedures were performed according to the protocols approved by the Jackson Laboratory and Columbia University’s Institutional Animal Care and Use Committee.

## Author contributions

S.W.M.J. conceived the project; C.Z., J.M.H., and S.W.M.J. designed the research; C.Z., H.L., Q.W., M.S., J.M.H., and S.W.M.J. performed research and analyzed data; C.M. managed mouse colony; C.Z., M.S., and S.W.M.J. wrote the manuscript. All authors read and edited the manuscript.

## Acknowledgments

The authors would like to thank Amy Bell for measuring intraocular pressure in mice, and Pete Finger for optic nerve sectioning and PPD staining.

## References

1. Adornetto A, Russo R, Parisi V. Neuroinflammation as a target for glaucoma therapy. Neural Regen Res. Mar 2019;14(3):391–394. doi:10.4103/1673-5374.245465

2. Russo R, Varano GP, Adornetto A, et al. Retinal ganglion cell death in glaucoma: Exploring the role of neuroinflammation. Eur J Pharmacol. Sep 15 2016;787:134–42. doi:10.1016/j.ejphar.2016.03.064

3. Shestopalov VI, Spurlock M, Gramlich OW, Kuehn MH. Immune Responses in the Glaucomatous Retina: Regulation and Dynamics. Cells. Aug 3 2021;10(8) doi:10.3390/cells10081973

4. Williams PA, Braine CE, Kizhatil K, et al. Inhibition of monocyte-like cell extravasation protects from neurodegeneration in DBA/2J glaucoma. Mol Neurodegener. Jan 22 2019;14(1):6. doi:10.1186/s13024-018-0303-3

5. Baudouin C, Kolko M, Melik-Parsadaniantz S, Messmer EM. Inflammation in Glaucoma: From the back to the front of the eye, and beyond. Prog Retin Eye Res. Jul 2021;83:100916. doi:10.1016/j.preteyeres.2020.100916

6. Howell GR, Soto I, Zhu X, et al. Radiation treatment inhibits monocyte entry into the optic nerve head and prevents neuronal damage in a mouse model of glaucoma. J Clin Invest. Apr 2012;122(4):1246–61. doi:10.1172/JCI61135

7. Harder JM, Williams PA, Braine CE, et al. Complement peptide C3a receptor 1 promotes optic nerve degeneration in DBA/2J mice. J Neuroinflammation. Nov 11 2020;17(1):336. doi:10.1186/s12974-020-02011-z

8. Harder JM, Braine CE, Williams PA, et al. Early immune responses are independent of RGC dysfunction in glaucoma with complement component C3 being protective. Proc Natl Acad Sci U S A. May 9 2017;114(19):E3839–E3848. doi:10.1073/pnas.1608769114

9. Howell GR, Macalinao DG, Sousa GL, et al. Molecular clustering identifies complement and endothelin induction as early events in a mouse model of glaucoma. J Clin Invest. Apr 2011;121(4):1429–44. doi:10.1172/JCI44646

10. Williams PA, Braine CE, Foxworth NE, Cochran KE, John SWM. GlyCAM1 negatively regulates monocyte entry into the optic nerve head and contributes to radiation-based protection in glaucoma. J Neuroinflammation. Apr 26 2017;14(1):93. doi:10.1186/s12974-017-0868-8

11. Margeta MA, Lad EM, Proia AD. CD163+ macrophages infiltrate axon bundles of postmortem optic nerves with glaucoma. Graefes Arch Clin Exp Ophthalmol. Dec 2018;256(12):2449–2456. doi:10.1007/s00417-018-4081-y

12. Williams PA, Harder JM, Foxworth NE, et al. Vitamin B(3) modulates mitochondrial vulnerability and prevents glaucoma in aged mice. Science. Feb 17 2017;355(6326):756–760. doi:10.1126/science.aal0092

13. Williams PA, Harder JM, John SWM. Glaucoma as a Metabolic Optic Neuropathy: Making the Case for Nicotinamide Treatment in Glaucoma. J Glaucoma. Dec 2017;26(12):1161–1168. doi:10.1097/IJG.0000000000000767

14. Zhang ZQ, Xie Z, Chen SY, Zhang X. Mitochondrial dysfunction in glaucomatous degeneration. Int J Ophthalmol. 2023;16(5):811–823. doi:10.18240/ijo.2023.05.20

15. Harder JM, Guymer C, Wood JPM, et al. Disturbed glucose and pyruvate metabolism in glaucoma with neuroprotection by pyruvate or rapamycin. Proc Natl Acad Sci U S A. Dec 29 2020;117(52):33619–33627. doi:10.1073/pnas.2014213117

16. Leruez S, Marill A, Bresson T, et al. A Metabolomics Profiling of Glaucoma Points to Mitochondrial Dysfunction, Senescence, and Polyamines Deficiency. Invest Ophthalmol Vis Sci. Sep 4 2018;59(11):4355–4361. doi:10.1167/iovs.18-24938

17. Jassim AH, Inman DM, Mitchell CH. Crosstalk Between Dysfunctional Mitochondria and Inflammation in Glaucomatous Neurodegeneration. Front Pharmacol. 2021;12:699623. doi:10.3389/fphar.2021.699623

18. Casson RJ, Chidlow G, Crowston JG, Williams PA, Wood JPM. Retinal energy metabolism in health and glaucoma. Progress in Retinal and Eye Research. Mar 2021;81 doi:ARTN 100881 10.1016/j.preteyeres.2020.100881

19. Chrysostomou V, Rezania F, Trounce IA, Crowston JG. Oxidative stress and mitochondrial dysfunction in glaucoma. Curr Opin Pharmacol. Feb 2013;13(1):12–15. doi:10.1016/j.coph.2012.09.008

20. Tribble JR, Otmani A, Sun SS, et al. Nicotinamide provides neuroprotection in glaucoma by protecting against mitochondrial and metabolic dysfunction. Redox Biol. Jul 2021;43 doi:ARTN 101988 10.1016/j.redox.2021.101988

21. Williams PA, Casson RJ. Glycolysis and glucose metabolism as a target for bioenergetic and neuronal protection in glaucoma. Neural Regen Res. Aug 1 2024;19(8):1637–1638. doi:10.4103/1673-5374.389638

22. Williams PA, Harder JM, Cardozo BH, Foxworth NE, John SWM. Nicotinamide treatment robustly protects from inherited mouse glaucoma. Commun Integr Biol. 2018;11(1):e1356956. doi:10.1080/19420889.2017.1356956

23. Williams PA, Harder JM, Foxworth NE, Cardozo BH, Cochran KE, John SWM. Nicotinamide and WLD(S) Act Together to Prevent Neurodegeneration in Glaucoma. Front Neurosci. 2017;11:232. doi:10.3389/fnins.2017.00232

24. van Horssen J, van Schaik P, Witte M. Inflammation and mitochondrial dysfunction: A vicious circle in neurodegenerative disorders? Neurosci Lett. Sep 25 2019;710:132931. doi:10.1016/j.neulet.2017.06.050

25. Zheng Z, Yuan R, Song M, et al. The toll-like receptor 4-mediated signaling pathway is activated following optic nerve injury in mice. Brain Res. Dec 13 2012;1489:90–7. doi:10.1016/j.brainres.2012.10.014

26. Farooq M, Batool M, Kim MS, Choi S. Toll-Like Receptors as a Therapeutic Target in the Era of Immunotherapies. Front Cell Dev Biol. 2021;9:756315. doi:10.3389/fcell.2021.756315

27. Kuzmich NN, Sivak KV, Chubarev VN, Porozov YB, Savateeva-Lyubimova TN, Peri F. TLR4 Signaling Pathway Modulators as Potential Therapeutics in Inflammation and Sepsis. Vaccines (Basel). Oct 4 2017;5(4) doi:10.3390/vaccines5040034

28. Zeng F, Zheng J, Shen L, Herrera-Balandrano DD, Huang W, Sui Z. Physiological mechanisms of TLR4 in glucolipid metabolism regulation: Potential use in metabolic syndrome prevention. Nutr Metab Cardiovasc Dis. Jan 2023;33(1):38–46. doi:10.1016/j.numecd.2022.10.011

29. Katare PB, Bagul PK, Dinda AK, Banerjee SK. Toll-Like Receptor 4 Inhibition Improves Oxidative Stress and Mitochondrial Health in Isoproterenol-Induced Cardiac Hypertrophy in Rats. Front Immunol. 2017;8:719. doi:10.3389/fimmu.2017.00719

30. Balic JJ, Albargy H, Luu K, et al. STAT3 serine phosphorylation is required for TLR4 metabolic reprogramming and IL-1beta expression. Nat Commun. Jul 30 2020;11(1):3816. doi:10.1038/s41467-020-17669-5

31. Chi W, Li F, Chen H, et al. Caspase-8 promotes NLRP1/NLRP3 inflammasome activation and IL-1beta production in acute glaucoma. Proc Natl Acad Sci U S A. Jul 29 2014;111(30):11181–6. doi:10.1073/pnas.1402819111

32. Dvoriantchikova G, Barakat DJ, Hernandez E, Shestopalov VI, Ivanov D. Toll-like receptor 4 contributes to retinal ischemia/reperfusion injury. Mol Vis. Sep 30 2010;16:1907–12.

33. Allensworth JJ, Planck SR, Rosenbaum JT, Rosenzweig HL. Investigation of the differential potentials of TLR agonists to elicit uveitis in mice. J Leukoc Biol. Dec 2011;90(6):1159–66. doi:10.1189/jlb.0511249

34. Morzaev D, Nicholson JD, Caspi T, Weiss S, Hochhauser E, Goldenberg-Cohen N. Toll-like receptor-4 knockout mice are more resistant to optic nerve crush damage than wild-type mice. Clin Exp Ophthalmol. Sep-Oct 2015;43(7):655–65. doi:10.1111/ceo.12521

35. Nakano Y, Shimazawa M, Ojino K, et al. Toll-like receptor 4 inhibitor protects against retinal ganglion cell damage induced by optic nerve crush in mice. J Pharmacol Sci. Mar 2017;133(3):176–183. doi:10.1016/j.jphs.2017.02.012

36. Mzyk P, Hernandez H, Le T, Ramirez JR, McDowell CM. Toll-Like Receptor 4 Signaling in the Trabecular Meshwork. Front Cell Dev Biol. 2022;10:936115. doi:10.3389/fcell.2022.936115

37. Sharma TP, Curry S, McDowell CM. Effects of Toll-Like Receptor 4 Inhibition on Transforming Growth Factor-beta2 Signaling in the Human Trabecular Meshwork. J Ocul Pharmacol Ther. Apr 2020;36(3):170–178. doi:10.1089/jop.2019.0076

38. Geiduschek EK, Milne PD, Mzyk P, Mavlyutov TA, McDowell CM. TLR4 signaling modulates extracellular matrix production in the lamina cribrosa. Front Ophthalmol (Lausanne). 2022;2 doi:10.3389/fopht.2022.968381

39. Chaiwiang N, Poyomtip T. The association of toll-like receptor 4 gene polymorphisms with primary open angle glaucoma susceptibility: a meta-analysis. Biosci Rep. Apr 30 2019;39(4) doi:10.1042/BSR20190029

40. Janssen SF, Gorgels TG, van der Spek PJ, Jansonius NM, Bergen AA. In silico analysis of the molecular machinery underlying aqueous humor production: potential implications for glaucoma. J Clin Bioinforma. Oct 28 2013;3(1):21. doi:10.1186/2043-9113-3-21

41. Lin Z, Huang S, Sun J, Xie B, Zhong Y. Associations between TLR4 Polymorphisms and Open Angle Glaucoma: A Meta-Analysis. Biomed Res Int. 2019;2019:6707650. doi:10.1155/2019/6707650

42. Shibuya E, Meguro A, Ota M, et al. Association of Toll-like receptor 4 gene polymorphisms with normal tension glaucoma. Invest Ophthalmol Vis Sci. Oct 2008;49(10):4453–7. doi:10.1167/iovs.07-1575

43. Anderson MG, Smith RS, Hawes NL, et al. Mutations in genes encoding melanosomal proteins cause pigmentary glaucoma in DBA/2J mice. Nat Genet. Jan 2002;30(1):81–5. doi:10.1038/ng794

44. Lahola-Chomiak AA, Footz T, Nguyen-Phuoc K, et al. Non-Synonymous variants in premelanosome protein (PMEL) cause ocular pigment dispersion and pigmentary glaucoma. Hum Mol Genet. Apr 15 2019;28(8):1298–1311. doi:10.1093/hmg/ddy429

45. De Moraes CG, John SWM, Williams PA, Blumberg DM, Cioffi GA, Liebmann JM. Nicotinamide and Pyruvate for Neuroenhancement in Open-Angle Glaucoma: A Phase 2 Randomized Clinical Trial. JAMA Ophthalmol. Jan 1 2022;140(1):11–18. doi:10.1001/jamaophthalmol.2021.4576

46. Hui F, Tang J, Williams PA, et al. Improvement in inner retinal function in glaucoma with nicotinamide (vitamin B3) supplementation: A crossover randomized clinical trial. Clin Exp Ophthalmol. Sep 2020;48(7):903–914. doi:10.1111/ceo.13818

47. Poltorak A, He X, Smirnova I, et al. Defective LPS signaling in C3H/HeJ and C57BL/10ScCr mice: mutations in Tlr4 gene. Science. Dec 11 1998;282(5396):2085–8. doi:10.1126/science.282.5396.2085

48. Vogel SN, Hansen CT, Rosenstreich DL. Characterization of a congenitally LPS-resistant, athymic mouse strain. J Immunol. Feb 1979;122(2):619–22.

49. Tolman NG, Balasubramanian R, Macalinao DG, et al. Genetic background modifies vulnerability to glaucoma-related phenotypes in Lmx1b mutant mice. Dis Model Mech. Feb 19 2021;14(2) doi:10.1242/dmm.046953

50. Chang B, Smith RS, Hawes NL, et al. Interacting loci cause severe iris atrophy and glaucoma in DBA/2J mice. Nat Genet. Apr 1999;21(4):405–9. doi:10.1038/7741

51. John SW, Smith RS, Savinova OV, et al. Essential iris atrophy, pigment dispersion, and glaucoma in DBA/2J mice. Invest Ophthalmol Vis Sci. May 1998;39(6):951–62.

52. John SW, Hagaman JR, MacTaggart TE, Peng L, Smithes O. Intraocular pressure in inbred mouse strains. Invest Ophthalmol Vis Sci. Jan 1997;38(1):249–53.

53. Savinova OV, Sugiyama F, Martin JE, et al. Intraocular pressure in genetically distinct mice: an update and strain survey. BMC Genet. 2001;2:12. doi:10.1186/1471-2156-2-12

54. Anderson MG, Libby RT, Gould DB, Smith RS, John SW. High-dose radiation with bone marrow transfer prevents neurodegeneration in an inherited glaucoma. Proc Natl Acad Sci U S A. Mar 22 2005;102(12):4566–71. doi:10.1073/pnas.0407357102

55. Howell GR, Libby RT, Jakobs TC, et al. Axons of retinal ganglion cells are insulted in the optic nerve early in DBA/2J glaucoma. J Cell Biol. Dec 31 2007;179(7):1523–37. doi:10.1083/jcb.200706181

56. Smith RS, John SWM, Nishina PM, Sundberg JP. Systematic Evaluation of the Mouse Eye: Anatomy, Pathology, and Biomethods. CRC Press; 2002.

57. Libby RT, Li Y, Savinova OV, et al. Susceptibility to neurodegeneration in a glaucoma is modified by Bax gene dosage. PLoS Genet. Jul 2005;1(1):17–26. doi:10.1371/journal.pgen.0010004

58. Howell GR, Libby RT, Marchant JK, et al. Absence of glaucoma in DBA/2J mice homozygous for wild-type versions of Gpnmb and Tyrp1. BMC Genet. Jul 3 2007;8:45. doi:10.1186/1471-2156-8-45

59. Libby RT, Anderson MG, Pang IH, et al. Inherited glaucoma in DBA/2J mice: pertinent disease features for studying the neurodegeneration. Vis Neurosci. Sep-Oct 2005;22(5):637–48. doi:10.1017/S0952523805225130

60. Hirt J, Porter K, Dixon A, McKinnon S, Liton PB. Contribution of autophagy to ocular hypertension and neurodegeneration in the DBA/2J spontaneous glaucoma mouse model. Cell Death Discov. 2018;4:14. doi:10.1038/s41420-018-0077-y

61. Yang XL, van der Merwe Y, Sims J, et al. Age-related Changes in Eye, Brain and Visuomotor Behavior in the DBA/2J Mouse Model of Chronic Glaucoma. Sci Rep. Mar 15 2018;8(1):4643. doi:10.1038/s41598-018-22850-4

62. Donahue RJ, Fehrman RL, Gustafson JR, Nickells RW. BCLX(L) gene therapy moderates neuropathology in the DBA/2J mouse model of inherited glaucoma. Cell Death Dis. Aug 10 2021;12(8):781. doi:10.1038/s41419-021-04068-x

63. Syc-Mazurek SB, Fernandes KA, Libby RT. JUN is important for ocular hypertension-induced retinal ganglion cell degeneration. Cell Death Dis. Jul 20 2017;8(7):e2945. doi:10.1038/cddis.2017.338

64. Ahmad A, Crupi R, Campolo M, Genovese T, Esposito E, Cuzzocrea S. Absence of TLR4 reduces neurovascular unit and secondary inflammatory process after traumatic brain injury in mice. PLoS One. 2013;8(3):e57208. doi:10.1371/journal.pone.0057208

65. Lee HS, Hattori T, Park EY, Stevenson W, Chauhan SK, Dana R. Expression of toll-like receptor 4 contributes to corneal inflammation in experimental dry eye disease. Invest Ophthalmol Vis Sci. Aug 17 2012;53(9):5632–40. doi:10.1167/iovs.12-9547

66. Chang JH, McCluskey PJ, Wakefield D. Toll-like receptors in ocular immunity and the immunopathogenesis of inflammatory eye disease. Br J Ophthalmol. Jan 2006;90(1):103–8. doi:10.1136/bjo.2005.072686

67. Titi-Lartey O, Mohammed I, Amoaku WM. Toll-Like Receptor Signalling Pathways and the Pathogenesis of Retinal Diseases. Front Ophthalmol. 2022;2 doi:10.3389/fopht.2022.850394

68. Duarte JN. Neuroinflammatory Mechanisms of Mitochondrial Dysfunction and Neurodegeneration in Glaucoma. J Ophthalmol. 2021;2021:4581909. doi:10.1155/2021/4581909

69. Ju WK, Perkins GA, Kim KY, Bastola T, Choi WY, Choi SH. Glaucomatous optic neuropathy: Mitochondrial dynamics, dysfunction and protection in retinal ganglion cells. Prog Retin Eye Res. Jul 2023;95:101136. doi:10.1016/j.preteyeres.2022.101136

70. Lee S, Van Bergen NJ, Kong GY, et al. Mitochondrial dysfunction in glaucoma and emerging bioenergetic therapies. Exp Eye Res. Aug 2011;93(2):204–12. doi:10.1016/j.exer.2010.07.015

71. Kalesnykas G, Oglesby EN, Zack DJ, et al. Retinal ganglion cell morphology after optic nerve crush and experimental glaucoma. Invest Ophthalmol Vis Sci. Jun 22 2012;53(7):3847–57. doi:10.1167/iovs.12-9712

72. Wang J, Struebing FL, Geisert EE. Commonalities of optic nerve injury and glaucoma-induced neurodegeneration: Insights from transcriptome-wide studies. Exp Eye Res. Jun 2021;207:108571. doi:10.1016/j.exer.2021.108571

73. Imai Y, Kuba K, Neely GG, et al. Identification of oxidative stress and Toll-like receptor 4 signaling as a key pathway of acute lung injury. Cell. Apr 18 2008;133(2):235–49. doi:10.1016/j.cell.2008.02.043

74. Morales-Nebreda L, Mutlu GM, Scott Budinger GR, Radigan KA. Loss of TLR4 does not prevent influenza A-induced mortality. Am J Respir Crit Care Med. May 15 2014;189(10):1280–1. doi:10.1164/rccm.201401-0193LE

75. Shirey KA, Blanco JCG, Vogel SN. Targeting TLR4 Signaling to Blunt Viral-Mediated Acute Lung Injury. Front Immunol. 2021;12:705080. doi:10.3389/fimmu.2021.705080

76. Rocha DM, Caldas AP, Oliveira LL, Bressan J, Hermsdorff HH. Saturated fatty acids trigger TLR4-mediated inflammatory response. Atherosclerosis. Jan 2016;244:211–5. doi:10.1016/j.atherosclerosis.2015.11.015

77. Rogero MM, Calder PC. Obesity, Inflammation, Toll-Like Receptor 4 and Fatty Acids. Nutrients. Mar 30 2018;10(4) doi:10.3390/nu10040432

